# Biocompatibility of KAPs-Depleted Residual Hair

**DOI:** 10.1101/2024.04.29.591537

**Authors:** Allison Meer, Aidan Mathews, Mariana Cabral, Andrew Tarabokija, Evan Carroll, Henna Chaudhry, Michelle Paszek, Nancy Radecker, Thomas Palaia, Roche C. de Guzman

## Abstract

This work is an in-depth investigation of the *in vitro* and *in vivo* biocompatibility of processed and treated residual human hair samples with intact cuticle layers. The specimens included oxidized hair with no melanin (BLH) and hair with medium-(M-KAP) and low-(L-KAP) levels of keratin associated proteins (KAPs), confirmed through gel electrophoresis, electron microscopy, and trichrome histological staining, in comparison to the untreated regular hair (REG) control. All hair groups, high KAPs (H-KAPs: REG and BLH), M-KAP, and L-KAP, were found to be non-cytotoxic in the adipose fibroblast cell line’s response to their extracts based on the ISO 10993-5 medical device biomaterial testing standard. *In vivo* mouse subcutaneous implantation (ISO 10993-6, local effects) at 2 weeks showed that the samples caused a foreign body response (FBR) with a thin fibrous encapsulation at a mean value of 28% relative skin dermis thickness; but notably, the L-KAP implant mitigated a statistically significant decrease in FBR area compared to H-KAP’s (REG and/or BLH) and a lower number of cells, including immune cells of mostly macrophages and mast cells on the biomaterial’s surface, normalized to implant and tissue coverage. In the bulk of the capsules, blood vessels and collagen extracellular matrix densities were similar among groups. These findings suggest that small globular KAPs diffuse out of the cortex to the host-biomaterial interface which induce a slightly elevated FBR but limited to the implant’s surface vicinity. On-going follow-up research focuses on purer keratin-based macromolecularly organized residual hair biomaterials, those with depleted KAPs, for drug-delivery gel implants as they are deemed the most biocompatible.

**Statement of Significance:** Human hair is an abundant biological product that is regularly discarded and wasted but has the potential to be a clinical implantable allograft biomaterial. There are currently just two FDA class II-510(k)-approved medical devices from hair, limited to surface / skin wound care use, and no class III-PMA or biologics-BLA implants. Also, these products and those in research and development phases are based on soluble keratin and KAPs extracts utilizing tedious processing conditions and requiring oxidation reaction for reassembly into gels and scaffolds. Here we describe that the insoluble residual hair biomaterials with organized keratin structure, higher-degree of disulfide crosslinks, and particularly those with depleted KAPs have increased biocompatibility based on pre-clinical ISO 10993 standards. This novel natural biomaterials are now being developed as drug-delivery implantable gels for clinical applications.

## 1. Introduction

The human hair fiber is an under-utilized biological product (biologic) that holds significant importance and potential for widespread adoption in the field of clinical medicine. Its structure is well-characterized: with a thin multi-layered and scaly surface, the cuticle (approximately 10% of the hair mass), and the majority of its bulk, the cortex (occupying 86%-90%). The hair’s main constituents are proteins, accounting to 98% by mass; most of which (at 85% of the total protein) are the cytoskeletal proteins: hair keratins and keratin associated proteins (KAPs) of about 80% and 5% (implied) weight fraction of the total dried hair, respectively (or ∼ 16:1 = keratins/KAPs mass ratio) [1–3]. Within the cortex, rod-shaped keratins are macromolecularly assembled into long strands of intermediate filaments (IFs, previously called microfibrils), with 7-10 nm in diameter. IFs are then bundled together as a 150-500 nm-wide macrofibril, with spaces filled-up and tightly-crosslinked by small globular matrix proteins, KAPs, which account for ∼ 40% of the macrofibril volume [1, 4]. The organized interaction of keratins and KAPs provides the high tensile strength of hair that is comparable to steel at normalized densities, and a greater elastic modulus than the collagen extracellular matrix (ECM) protein [5, 6].

Biomedical studies of hair are based on the soluble and extractable fractions of these combined keratins and KAPs (which are challenging to separate) in their reorganized scaffold and gel formulations of reduced and/or oxidatively-modified versions, often referred to and simplified as “keratin biomaterials.” Our research group and others have demonstrated their biocompatibility and drug-delivery activities in animal models for tissue engineering and regenerative medicine [7–12]. Despite numerous promising experimental and pre-clinical outcomes, only very few keratin biomaterial products have undergone clinical trials/studies (IDE), U.S. Food and Drug Administration (FDA) 510(k) premarket notification clearances, and commercialization; however, applications were limited to skin treatments as wound care dressings (ProgenaMatrix from human hair by ProgenaCare, Marietta, GA, and Keratec / Keraderm / Keramatrix from sheep wool by Keratec, Christchurch, New Zealand and Molecular Biologicals, Charlottesville, VA) and as a topical cream (KeraStat from human hair by KeraNetics, Winston-Salem, NC) [13–18]. Currently, there are still no FDA-approved and marketed class III medical devices (PMA), biologics (BLA route), or combination products employing keratin biomaterials. Some of the scientific reasons include the technical complexity, heterogeneity, and loss of mechanical integrity during keratins and KAPs reassembly and material preparation.

Alternatively, the insoluble left-over residual hair samples after chemical processing can address the abovementioned issues, and thus may be better candidate biomaterial fibers for clinical translation. They are simpler to make and keep hair keratins and KAPs, but this time in natively organized structures with higher amounts of disulfide linkages. Other important proteins including histones with antimicrobial activity [19] may also be retained. However, the insoluble hair remnants are just discarded, largely ignored, and unstudied. Accordingly, aside from investigating the biocompatibility of residual hairs to get their important safety information, which is the paper’s primary objective, we want to know if certain hair components elucidate differences in responses. Specifically, we tested REG = regular untreated black/dark hair serving as a control, and three groups of residual hairs (with preserved cuticles to minimize absorption and degradation) based on processing conditions leading to the removal of melanin (as BLH = bleached hair) and degrees of KAPs depletion (as M-KAP = medium-level KAPs and L-KAP = low-level KAPs) for ISO 10993 – Biological evaluation of medical devices: *in vitro* cytotoxicity and *in vivo* local effects after implantation, with results serving as pre-clinical data for eventual FDA approval. We emphasized the microscopic image analysis of the fibrous capsule foreign body response against the subcutaneous implantation of these hair biomaterials.

## 2. Materials and methods

### 2.1 Preparation and characterization of residual hair biomaterials

*REG, BLH, M-KAP, and L-KAP* Hair samples were donated by and collected from local salons (Long Island, NY). After being washed with soap to remove dirt, the samples were dehydrated in ethanol (EtOH) and delipidized in 2:1 volumes (V) of chloroform : methanol at 37 °C, overnight (O/N) with shaking. This hair was set aside as untreated regular hair control (**REG**). Three treatment groups of residual hairs were prepared, namely bleached hair (**BLH**), hair containing medium-level KAPs (**M-KAP**), and hair containing low-level KAPs (**L-KAP**). <u>For BLH</u> – REG’s surfaces were applied with a bleaching mix (1 V bleach powder (Perfect Blond, Blond Forte, Medley, FL) to 2 V developer (Crème 40 V, Clairol Professional, Petit-Lancy, Geneva, Switzerland)) wrapped in a foil for 40 min and rinsed with water, repeated twice. <u>For M-KAP</u> – REG was incubated at 50 mg/mL in a solution containing 25 mM tris, 25% (V/V) EtOH, 25 mM dithiothreitol (DTT), 350 mM β-mercaptoethanol (BME), 8 M urea, at pH 9.5 for 4 d at 37 °C with shaking and residual hair washed and rinsed with water. A no-EtOH control sample was included for comparison. <u>For L-KAP</u> – REG was instead treated with 2.15 M tris, 25% EtOH, and 1 M thioglycolic acid (TGA) at pH 9 and extensively rinsed in water, repeated up to three cycles (the L-KAP implant utilized 2 cycles).

#### Protein content

Soluble extracts from hair treatment processes were collected, dialyzed against water using a 3.5-kDa molecular weight cut-off (MWCO), and quantified for total protein using DC Assay (Bio-Rad, Hercules, CA) and melanin at 432-nm absorbance (brown, based on spectral scan) with a synthetic melanin reference (Sigma-Aldrich, St. Louis, MO). Proteins were visualized via sodium dodecyl sulfate-polyacrylamide gel electrophoresis (SDS-PAGE) with Precision Plus Protein Dual Color Standards (Bio-Rad) at volumes normalized to the initial hair mass. Additionally, REG and BLH residual hairs were subjected to 0.5 M TGA, pH 11 reduction followed by 100 mM tris solubilization and fractions 1 and 2, respectively, were quantified for melanin content and run in SDS-PAGE to determine their protein contents.

#### Scanning electron microscopy (SEM)

Samples were dehydrated in increasing EtOH (70%, 95%, and 100% for 5 min each). After surface sputter coating (EMS-550 Sputter Coater, Electron Microscopy Sciences, Hatfield, PA) with gold to provide electron conductivity, specimens were imaged using a Q250 SEM scanning electron microscope (FEI, Hillsboro, OR) at 30 kV for biomaterials’ surface visualization.

#### Transmission electron microscopy (TEM)

Hair samples were fixed in 4% glutaraldehyde buffered in 0.1 M sodium cacodylate buffer, pH 7.5, washed in sodium cacodylate buffer, post-fixed in buffered 1% osmium tetroxide, en bloc stained with a saturated solution of uranyl acetate in 40% EtOH, dehydrated in a graded series of ethanol, infiltrated with Epon epoxy resin (LADD LX112, Ladd industries, Burlington, VT), and embedded. The blocks were sectioned with a Reichert Ultracut microtome at 70-nm. The resulting grids were then post-stained with a 1% aqueous uranyl acetate followed by 0.5% aqueous lead citrate and scoped on a Zeiss EM 900 transmission electron microscope retrofitted with an SIA L3C digital camera (SIA, Duluth, GA) to image hairs’ internal micro and ultrastructure.

### 2.2 In vitro cell culture with hair samples and their extracts

The *in vitro* cytotoxicity or biocompatibility of treated hairs were evaluated according to ISO 10993-5: Biological evaluation of medical devices, Tests for *in vitro* cytotoxicity, Annex C: MTT assay [20] through direct contact and extract tests. L-929 mouse adipose areolar subcutaneous fibroblasts line (NCTC clone 929, ATCC, Manassas, VA) were cultured in Dulbecco’s Modified Eagle’s Medium (DMEM) with 10% (V/V) fetal bovine serum and 1× antibiotic-antimycotic agent in a 96-well plate at 100-mL with 10^4^ cells/well for 24 h in a mammalian cell incubator (37 °C, 5% CO_2_, >90% relative humidity). <u>For the direct contact</u> – Hair samples (REG, BLH, M-KAP, and L-KAP), rinsed in 70% EtOH then water (to sterilize), were placed directly on top of the cell monolayer covering ∼ 50% area. <u>For the extract testing</u> – Specimens were surfaced-sterilized then soaked in culture medium (at 3 cm^2^/mL material surface area to medium volume ratio) and incubated for 3 d at 37 °C with shaking for extraction of any leachable substances, in accordance to ISO 10993-12: Sample preparation and reference materials [21], including latex (and 0.6% dimethyl sulfoxide added to cells) and polypropylene, positive (+) and negative (–) biomaterial cytotoxicity controls, respectively. 100-mL of these extracts at different dilutions (1, 0.5, 0.25, 0.125, and 0 × blank) were added to cultured cells. <u>For both</u> – Cells at direct contact with hair or their extracts were incubated for 24 h to induce their effects, followed by addition of 50 mL of 1 mg/mL (3-(4,5-dimethylthiazol-2-yl)-2,5-diphenyltetrazolium bromide) (MTT) in DMEM for 2 h, and finally medium was removed, formazan crystals dissolved with isopropanol, and absorbance read at 570-nm. Experimental groups were compared to + and – cytotoxicity controls.

### 2.3 Animal implantation for biocompatibility assessment of hair biomaterials

This short-term animal study was supported by the approved Hofstra Institutional Animal Care and Use Committee (IACUC) protocol, following ISO 10993-2: Animal welfare requirements [22] and ISO 10993-6: Tests for local effects after implantation [23]. Male, adult (11-w) CD-1^®^ IGS outbred albino mice (Charles River, Wilmington, MA) were anesthetized using isoflurane delivered via the SomnoSuite low-flow system (Kent Scientific, Torrington, CT) and subcutaneously implanted (into the subcutaneous white adipose tissue (sWAT) or fat tissue layer) on their dorsal back region with a surface-sterilized and water-rinsed control regular hair (**REG**) and treated hair groups: bleached hair (**BLH**) and medium (**M-KAP**) and low KAPs-containing (**L-KAP**) hairs (∼ 0.1 mL, cut at < 5-mm, ≥ 80 strands) at n = 4 mice (biological replicates) each. At 2-w post-operative (PO), mice were euthanized and macroscopically evaluated for signs of local inflammation, toxicity, and adverse effects. Their dorsal skin was exposed, and implants were recovered together with their surrounding fibrous capsules, fixed in 10% neutral buffered formalin for 3 d, and stored in 70% EtOH.

### 2.4 Histology and image analysis

All fixed samples (fibrous tissues with encapsulated hair implants) were trimmed, processed, and embedded in paraffin blocks (Paraplast X-TRA, Sigma-Aldrich) based on the protocol from our recent research paper [24]. After microtome sectioning at a thickness of 5-mm and rehydration, samples were stained with Masson’s trichrome (MT, connective tissue stain, ab150686 kit, Abcam, Waltham, MA) to highlight and detect connective tissue and immune cell components (black/dark for cell nuclei, red for cytoskeleton and cytosol of cells, and blue for the collagen extracellular matrix (ECM)), dehydrated, mounted using Histomount (Thermo Fisher Scientific, Waltham, MA), and imaged using Cytation 5 Cell Imaging Multimode Reader (Agilent, Santa Clara, CA), and Primostar 3 with Excelis (Zeiss, White Plains, NY) at low and high magnifications. Image processing and analysis were performed in GIMP (gimp.org), ImageJ (NIH, Bethesda, MD), MATLAB (MathWorks, Natick, MA), and Excel (Microsoft, Redmond, WA).

#### Fibrous encapsulation

The foreign body response (FBR) fibrosis due to hair implants was quantified through measurement of fibrous tissue wall thickness (outermost implant to outermost capsule) and area occupied by the fibrosis relative to implant area excluding internal airgaps due to processing (fibrous to implant area ratio). Additionally, hair implant thicknesses were obtained as the narrowest region in implants were cut in cross and longitudinal sections. The number of blood vessels in the FBR tissue based on morphology was manually counted per fibrous tissue area.

#### FBR cells and ECM evaluation

The amount of cells in the FBR was reported as cells’ area in tissue = cells/(cells + ECM) area ratio. Tissue definition was simplified as “cells + ECM”, where cells as red versus ECM as blue signals. The light background was converted to black of colored images including implant areas, split into red, green, and blue channels, and the mean of the intensities (ranging from 0 = no signal to 255 = full signal) of red and blue channels was quantified. The normalized ECM density (density/area) was also computed using MATLAB. Blue-channel images were read using imread function. The ECM area and density were measured with sum of logical (0 as absent and > 0 as present) and mean (average of signal intensities) functions, respectively.

#### Semi-quantitative scoring for local implantation effects

Cell types and abundance at high-powered field of view (at 40× objective lens for polymorphonuclear cells including mast cells, lymphocytes, plasma cells, macrophages, giant cells, and necrotic cells) and biological responses (neovascularization, fibrosis, and fatty infiltrate) were scored 0-4 and summed according to ISO 10993-6, Annex E: Examples of evaluation of local biological effects after implantation [23]. Average scores were subtracted by a reference score of 2.9 (considered the limit for no adverse effect) and categorized as minimal or no reaction for 0-2.9, slight for 3-8.9, moderate for 9-15, and severe for ≥ 15.1 mean scores.

### 2.5 Statistical methods

Images were processed in GIMP and PowerPoint (Microsoft). Data and graphs were evaluated and generated in Excel and MATLAB. Technical replicate values were averaged, and biological replicates’ values were again averaged then reported as mean (AVE) ± 1 standard deviation (STD). Student’s t-test and analysis of variance (ANOVA) with Tukey-Kramer were used for multiple comparison using a probability (p) < 0.05 deemed significantly different and represented as: ***p < 0.001, **p < 0.01, and *p < 0.05.

## 3. Results

### 3.1 Residual hair implant properties

The treated residual hairs were visibly distinguishable (**Fig. 1A**) from the untreated, regular hair (REG) control. Bleached hair (BLH) appeared off-white to white, while M-KAP and L-KAP were still mostly dark with some lighter strands but exhibited a highly noticeable waviness and curvature. The final mass yields from the initial hair mass of BLH, M-KAP, and L-KAP were 48%, 65%, and 59%, respectively.

**Fig. 1.**
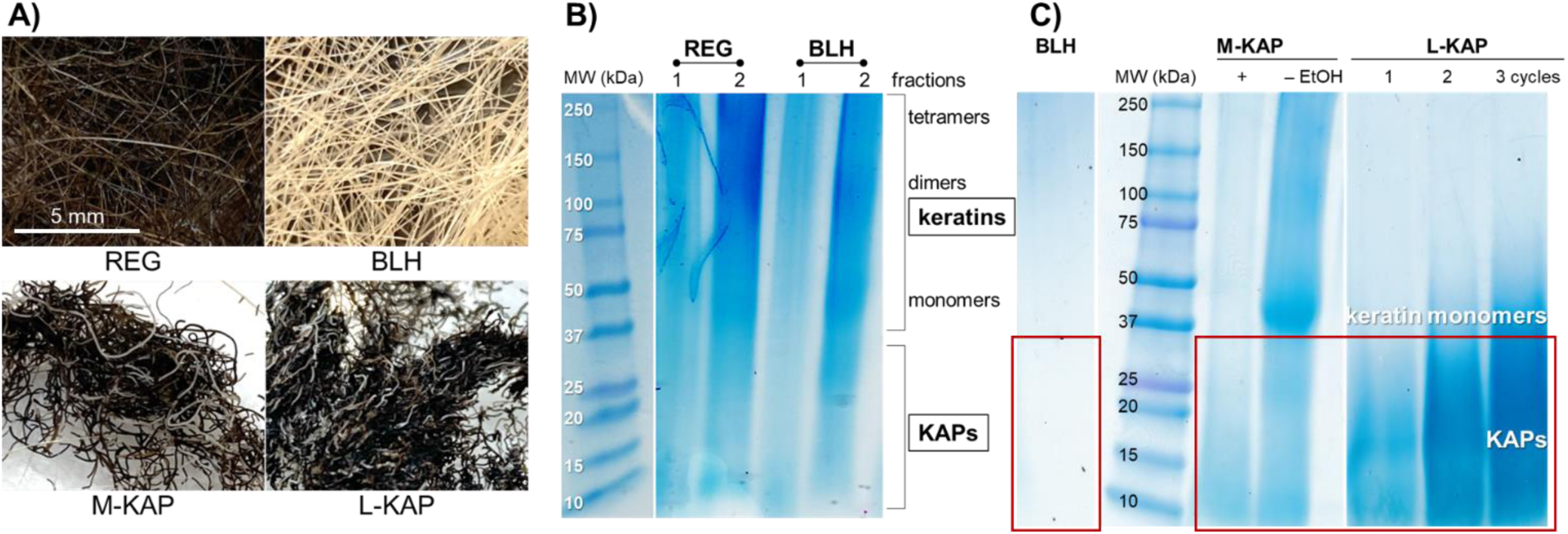
**A)** Untreated (REG = regular) and treated residual hair groups (BLH = bleached hair, M-KAP = medium KAPs, and L-KAP = low KAPs content). Gel electrophoresis of **B)** TGA and tris-solubilized fractions (1 and 2) of REG and BLH with high MWs representing keratins and low MWs for KAPs, and **C)** soluble proteins during the treatment processes of BLH, M-KAP, and L-KAP (boxed region highlight the KAPs removed during the residual hair preparation). In addition, removal of EtOH from the M-KAP’s extraction solution resulted in dissolution and release of keratins. Increase in extraction cycles using L-KAP’s medium released increasing amounts of low-MW KAPs.

BLH contained negligible (0.000) melanin compared to 0.023 mg/mL in REG extract (or 1:2174 relative to initial hair mass). TGA-reduced and tris-solubilized bleached hairs demonstrated that despite the differences in melanin, their hair protein composition of mostly keratins and KAPs were similar to REG (**Fig. 1B**). Protein bands in the low MWs (representing KAPs) and higher MWs: 40-60 kDa monomers, 80-120 kDa heterodimers, and 160-240 kDa tetramers of hair keratins exhibited general similarities in patterns and concentrations for both REG and BLH.

#### L-KAP residual hair has depleted amounts of KAPs

During the preparation of residual hair, soluble materials removed displayed almost no protein bands for BLH, but with relatively lower amounts in M-KAP and higher amounts in L-KAP of the smaller KAP proteins (**Fig. 1C**: boxed regions), implying that REG and BLH contained the highest quantity (H-KAP), then M-KAP with a medium-level, and finally L-KAP with the lowest amounts of keratin associated proteins. Ethanol in the extraction solution prevented the release of high MW keratins, which were retained in the residual hairs (while removal (-) of EtOH made keratins solubilized, **Fig. 1C**: M-KAP, - EtOH). Additional extraction cycle in L-KAP preparation released the highest KAPs amount but possibly included monomeric keratins (**Fig. 1C**: L-KAP, 3 cycles). It is noted that the L-KAP implant utilized 2 cycles, which preserved more keratins.

Electromicrographs of the materials’ topography (**Fig. 2**) and cross-section (**Fig. 3**) verified the following specific hair treatment and processing effects. The cuticle structure (thin, at < 5 mm, scaly outermost region observed with 5-7 layers in REG) was preserved in all treatments. But compared to the regular hair, BLH’s cuticle appeared irregular and smoother with less layers at 3-4 (**Fig. 3**: BLH), while M-KAP and L-KAP’s were wrinkled due to less cortical contents in these two groups (**Fig. 2**). Melanin granules or melanosomes (at submicron in diameter), which are typically found in the cortex (thick, inner bulk), were absent in the bleached residual hair (**Fig. 3**: BLH), leaving circular empty spaces behind. In M-KAP and L-KAP hairs, some melanosomes were also missing and distorted in shape. Distinct and organized macrofibrils composed of keratins and KAPs were hard to observe on their exposed cortex among all groups. In M-KAP and L-KAP’s TEM, loose cortical matrices were evident with lighter contrast or shade in M-KAP’s cortex versus REG’s and were especially highlighted in L-KAP where gaps that used to be filled with globular keratin associated proteins were clearly observed (**Fig. 3**: L-KAP, arrow). The decreased matrix of KAPs, compared to REG and BLH, contributed to their shriveled configuration (**Fig. 2**: M-KAP and L-KAP, main and insets).

**Fig. 2.**
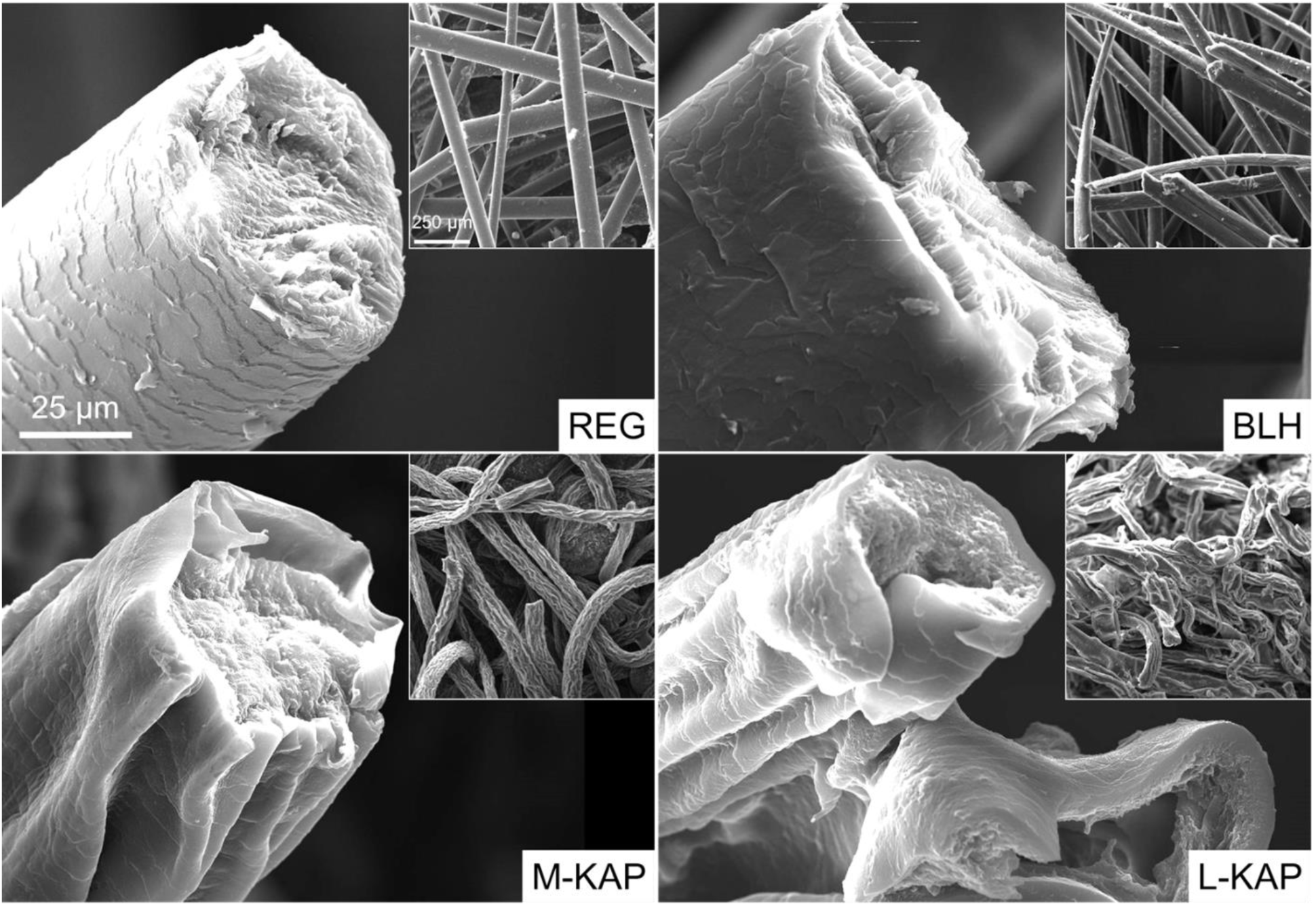
SEM images displaying representative hair strands with their outer cuticle layers and exposed inner cortical bulk. (Inset) Lower magnification of bundles of hair. M-KAP and L-KAP have assumed shriveled structures.

**Fig. 3.**
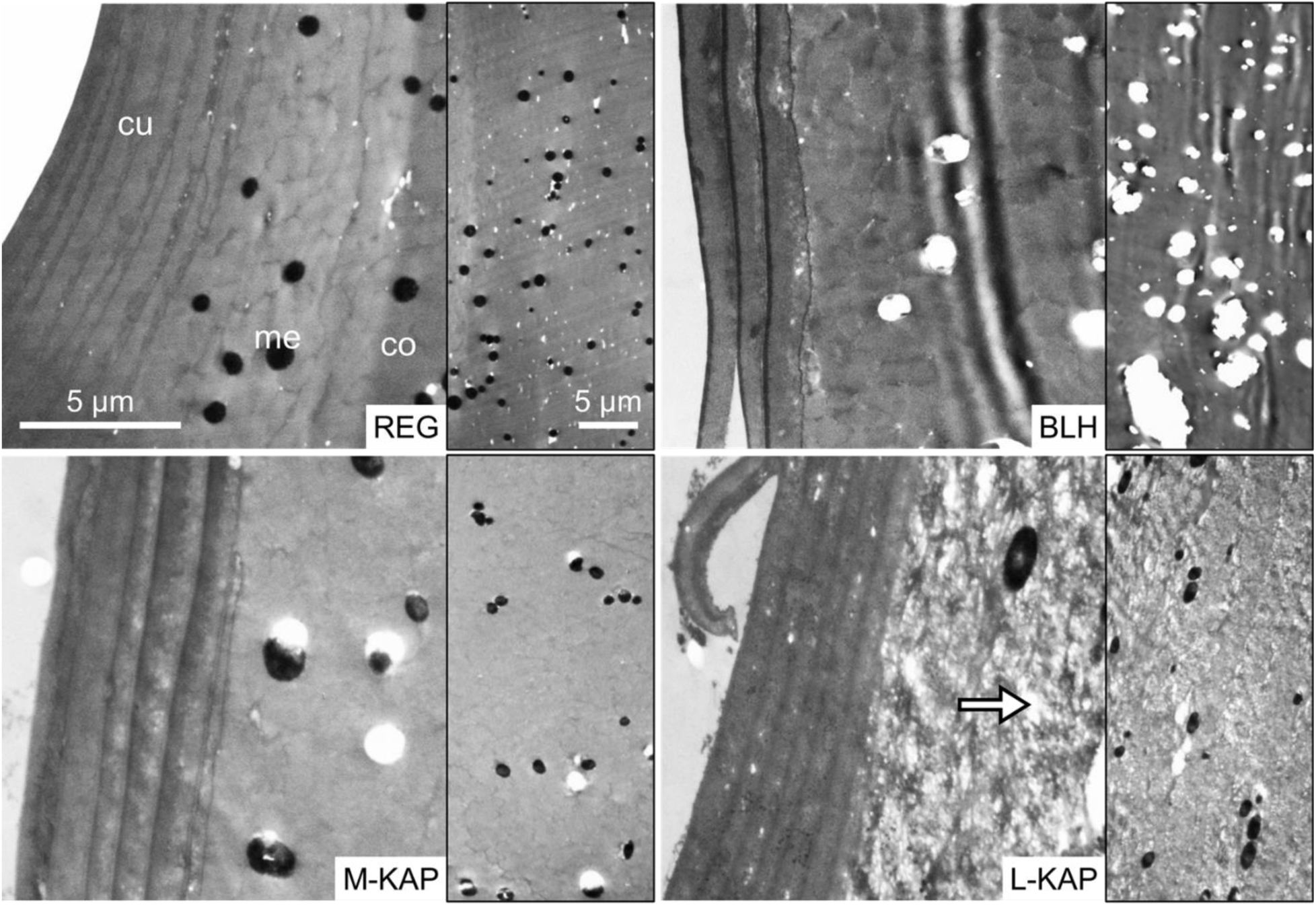
TEM of cross-sections of hair samples with notable structures: cu = cuticle, co = cortex, and me = melanin granules. The arrow in L-KAP are gaps used to be occupied by KAPs. (Right bordered section) Lower magnification of the cortical region with absence of melanin in the BLH group and looser matrices in M-KAP and L-KAP.

The summary of hair biomaterial properties is tabulated in **Table 1**.

**Table 1.** Physical properties of processed and treated hair biomaterials and analysis leading to conclude the relative abundance of keratin associated proteins (KAPs) content. SEM = scanning electron microscopy, TEM = transmission electron microscopy, MT = Masson’s trichrome staining, NA = not applicable, since unprocessed.

| Hair groups | Acronym meaning | Processing condition | Small protein solubles in processing<br>Fig. 1C | Macro color and appearance<br>Fig. 1A | SEM surface shape<br>Fig. 2 | TEM of melanin in the cortex<br>Fig. 3 | TEM of the cortical matrix<br>Fig. 3 | MT histology color of fibers<br>Fig. 6E | KAPs content of hair relative to REG |
| --- | --- | --- | --- | --- | --- | --- | --- | --- | --- |
| H-KAP: REG | High-KAPs: Regular hair | untreated control | NA | black and very dark, straight | straight, intact cuticle and cortex | a lot | tight, dark contrast | yellowish | same, **high** |
| H-KAP: BLH | High-KAPs: Bleached hair | treated residual hair | very few | very light to off-white, straight | straight, intact cuticle | absent, leaving holes behind | mostly tight, dark contrast | mostly yellowish with some red | almost same, **high** |
| M-KAP | Medium-KAPs hair | treated residual hair | some | very dark with few light, wavy | wrinkled, intact cuticle | mostly present | slightly loose, lighter contrast | mostly red | lower, **medium** |
| L-KAP | Low-KAPs hair | treated residual hair | a lot | dark with few light, wavy | wrinkled, intact cuticle | mostly present | loose, regions of dark and light contrast | mostly red | depleted, **low** |

### 3.2 Cell culture biocompatibility response

The direct contact *in vitro* test after 1 day decreased the cultured fibroblast viability in the non-cytotoxic polypropylene (PP) control down to 9%, while in hair samples to 16% ± 7%, compared to the blank (cells only, no material contact). This is likely due to deprivation of nutrients and respiration, including limited access to dissolved oxygen, hence not considered an appropriate cytotoxicity testing model for this system. Still, comparing the fibroblast survival relative to the PP values showed statistical similarities (p ≥ 0.1705) among hair biomaterials (BLH, M-KAP, and L-KAP) (**Fig. 4A**), with slightly elevated averages but high variability (standard deviation) in both M-KAP and L-KAP. This result indicated that all groups (PP, REG, and treated hairs) have the same viability effect when directly in contact with cultured cells.

**Fig. 4.**
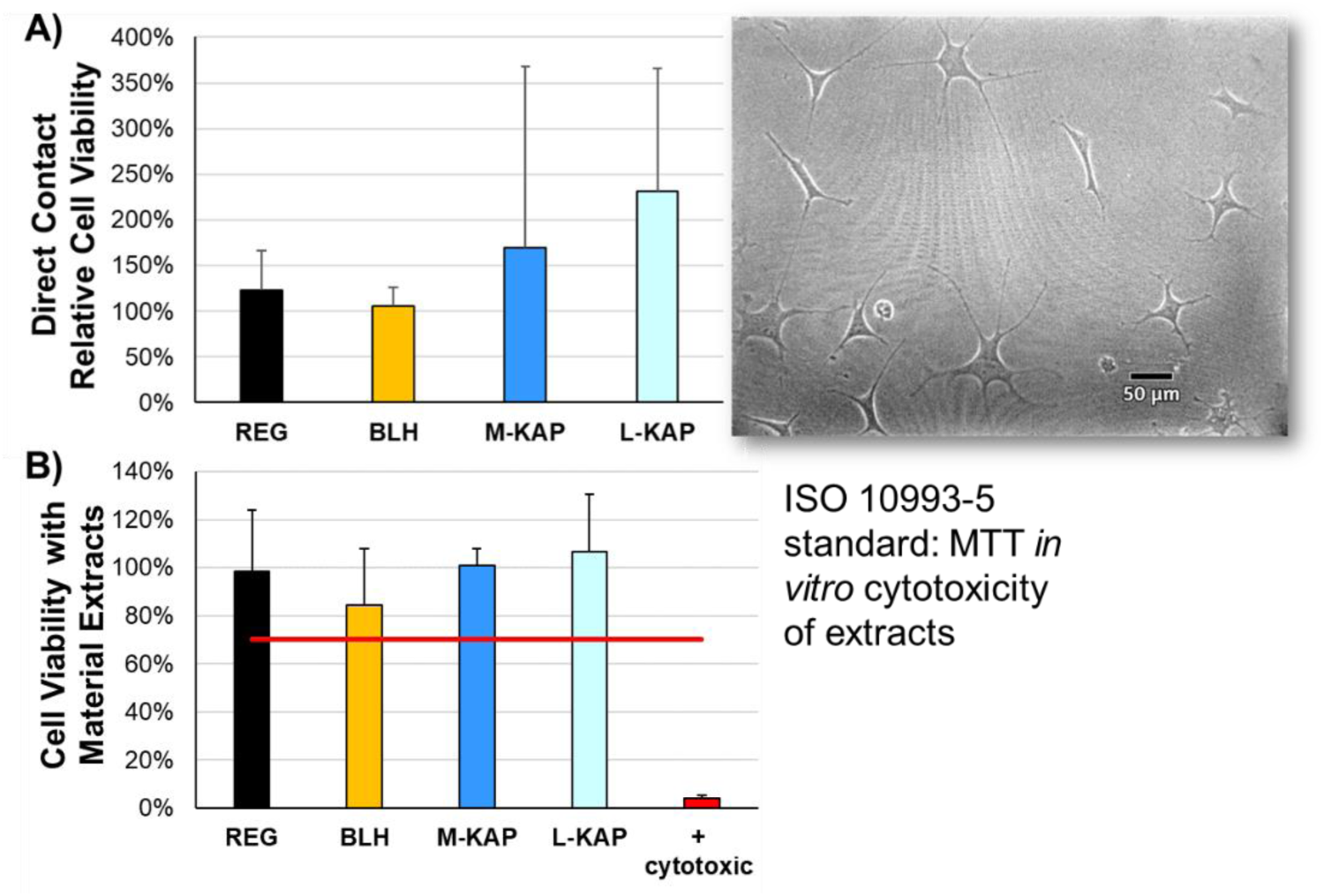
*In vitro* cytotoxicity testing with L-929 adipose fibroblast line through **A)** direct contact (showing the appearance of cells in phase contrast microscopy) and **B)** with extracts or leachables following the ISO 10993-5 MTT standard. No significant differences were observed among all hair groups and compared to the untreated and negative cytotoxicity controls, demonstrating *in vitro* biocompatibility and non-cytotoxicity. The viability values for all hair groups (REG, BLH, M-KAP, and L-KAP) were above the 70% cut-off (red line), thereby satisfying the ISO standard.

#### Hair samples are biocompatible, *in vitro*

Testing the hair extracts (instead of the prior direct contact method) is the ISO-recommended medical device’s biomaterial *in vitro* biocompatibility standard. MTT assay of hair extracts proved non-cytotoxicity as all viability values including different dilutions (1, 0.5, 0.25, and 0.125 ×) were > 70% compared to untreated (0 ×) control cells (**Fig. 4B**). This test confirmed that REG, BLH, M-KAP, and L-KAP residual hairs have no harmful water-leachable agents and pass the safety standard ISO 10993-5 as potential medical device biomaterials.

### 3.3 Mouse subcutaneous implantation biocompatibility response

#### Hair samples are biocompatible, *in vivo*

The dorsal skin flaps where hair implants were attached to, had completely-healed incision wounds (at 2 w, PO) with no observable hematoma and no gross inflammation (no redness and no edema) across all 4 groups (REG, BLH, M-KAP, and L-KAP) and among their biological replicates. No animals were compromised and lost during the study duration. Regional lymph nodes, particularly the axillary and inguinal [25], looked normal and were not swollen. The macroscopic fibrous capsule foreign body response (FBR) was generally small, thin, and almost clear, with encapsulated implants observable inside (**Fig. 5A**: representative of the L-KAP). The degree of vasculature appeared similar in regions with and without the implants and FBR tissues.

**Fig. 5.**
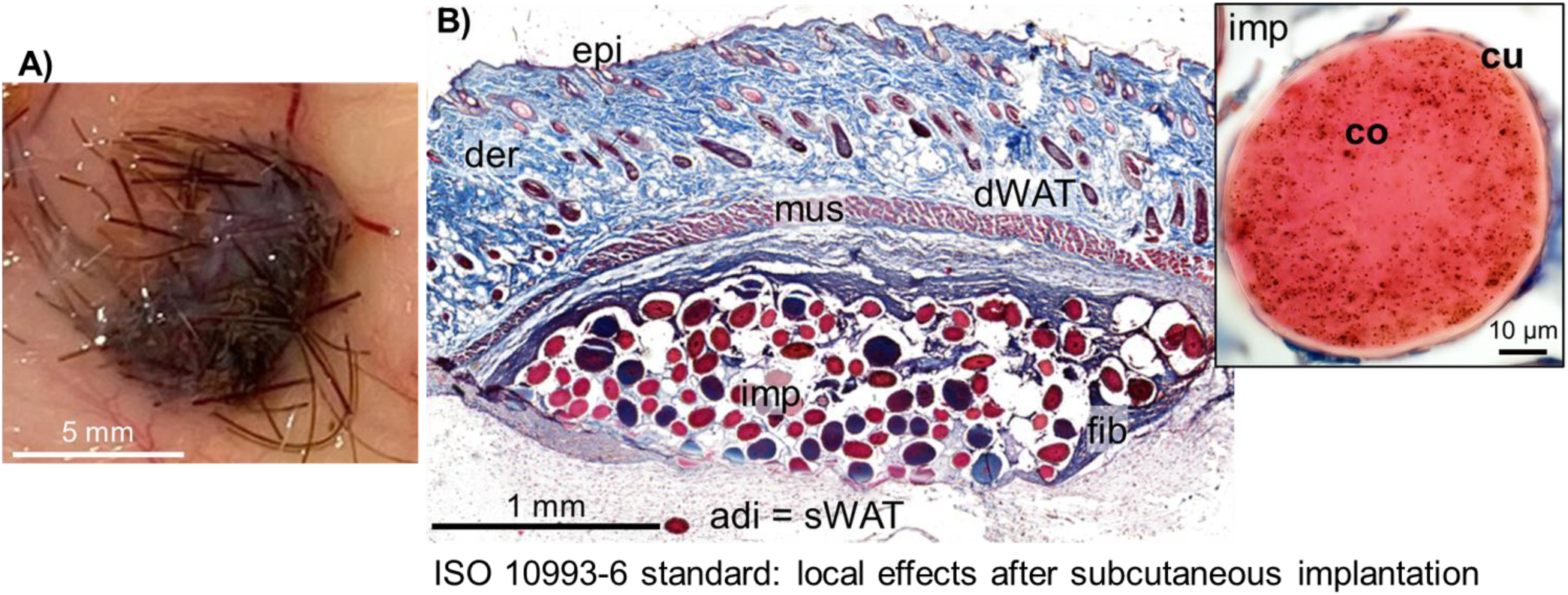
**A)** Inner dorsal skin flap showing an attached implant and its fibrous capsule response (almost clear) at 2 w, post-operative, and **B)** after Masson’s trichrome histology with key structural features: Epi = epidermis, der = dermis, and dWAT = dermal white adipose tissue of the skin (with mouse hairs), mus = muscle tissue (found in mouse, but not in humans), adi = sWAT = adipose or subcutaneous WAT, fib = fibrous capsule foreign body response, and imp = hair implants (within the fibrous capsule pocket) with a magnified example showing the clear cu = cuticle and reddish co = cortex regions (inset).

#### L-KAP induced the least fibrous capsule response

Regrettably, two samples were lost during the tissue processing, reducing the biological sample size of REG and M-KAP to n = 3. Histological staining with Masson’s trichrome (MT) of the rest generated good mouse skin cross-sections showing the skin layers: epidermis, dermis and dermal white adipose tissue (dWAT) [26, 27], the panniculus carnosus muscle [28], and subcutaneous WAT (sWAT) or fat, with the fibrous capsule FBR tissue found in between the muscle and sWAT layers. The hair implants can be observed encapsulated inside this fibrous tissue (**Fig. 5B**: L-KAP representative). They induced thin (at < 0.3 mm) fibrous capsules (**Fig. 6A**), specifically with 32% ± 17%, 43% ± 12%, 19% ± 8%, and 19% ± 5% for REG, BLH, M-KAP, and L-KAP, respectively, or an average 28% ± 11% of compared to the thickness of the skin dermis. Pairwise, both M-KAP and L-KAP’s FBR tissues were found to be statistically (*p = 0.033 and **p = 0.009) thinner than BLH’s. The areas occupied by the capsule (minus internal spaces) relative to implants were < 16-fold (**Fig. 6B**) with L-KAP generating statistically less (*p at least 0.044) than REG and BLH implants. The proportion of cells in the FBR tissue was discovered to be in accordance with the capsule thickness pattern, where more cells situated in BLH than in M-KAP and L-KAP (*p at least 0.0497) (**Fig. 6C**). On the other hand, the density (or thickness per area) of the collagen ECM secreted by fibroblasts of the capsule tissue was determined to be similar (p > 0.24) for all groups, at 1.8 ± 0.1 fold per mm^2^ (**Fig. 6D**). Both cells (as cell’s red MT-stained cytoplasm) and ECM (as blue MT-stained collagen) of the FBR and hair implants (within the tissue as rounded cross- and sometimes longitudinal-sections), and quantified measurement trends can be seen in selected regions of interest among sample replicates in **Fig. 6E**.

**Fig. 6.**
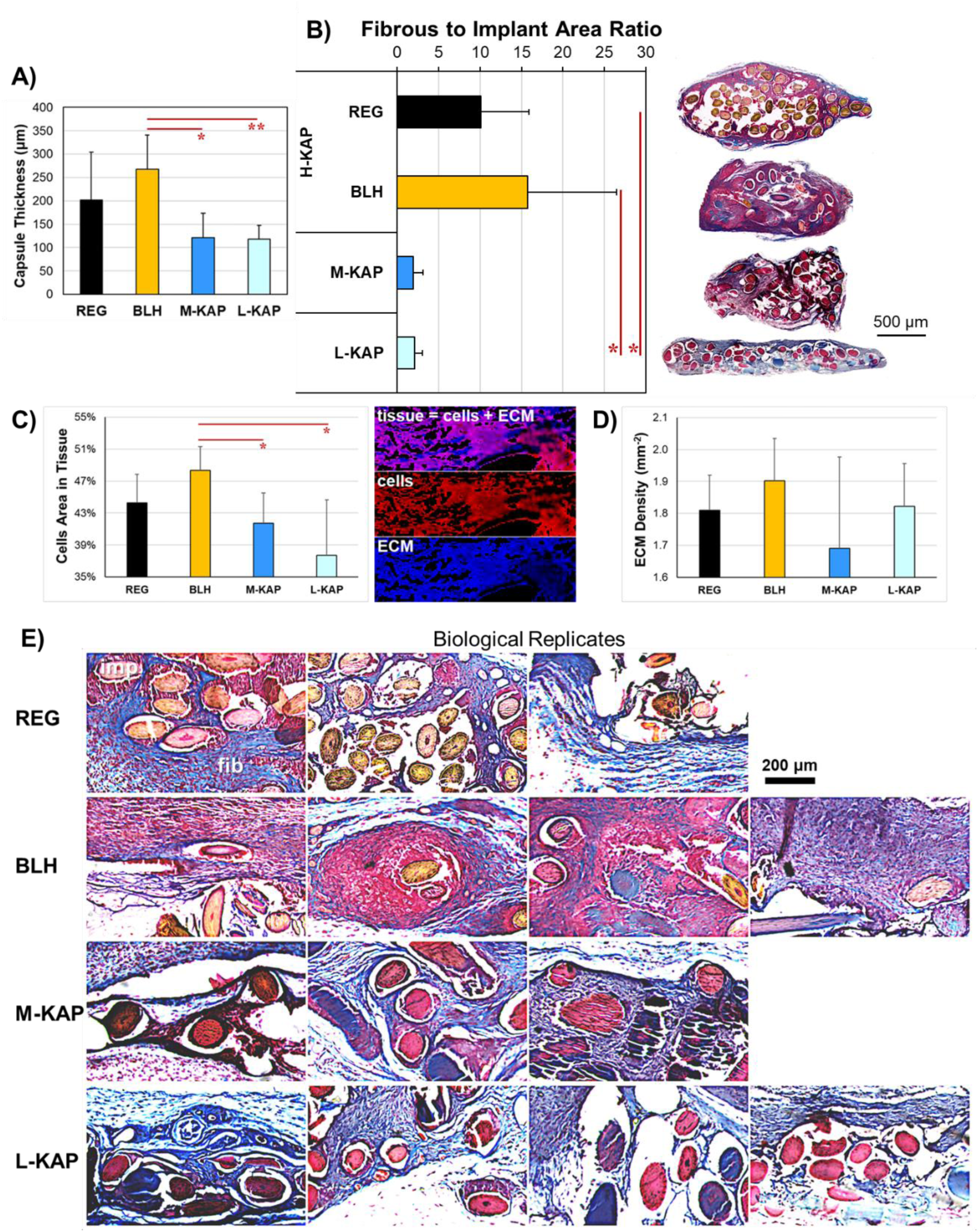
The fibrous capsule foreign body response to subcutaneous hair implants (REG, BLH, M-KAP, and L-KAP) showing their **A)** wall thicknesses, measured from the outermost implant to outermost fibrous tissue, are all relatively thin at < 0.3 mm, and **B)** areas (normalized to implants) at < 16-fold with group representative full-size histological sections (right). Simplified image analysis of the Masson’s trichrome staining denoted the tissue as cells = red plus ECM = blue signal in a black background resulting in **C)** area covered by the cells (cells/tissue) at < 49% and **D)** density of ECM (degree of blueness) normalized to tissue area at < 2-fold/mm^2^. **E)** Fibrous capsule (fib) and hair implant (imp) trichrome staining and morphologies among biological replicates of all groups. Fibrous tissues appear as blue, red, and blue-red combinations, surrounding the sectioned hair implants that stained as yellow, mostly for REG and BLH, while reddish for M-KAP and L-KAP.

#### Removal of KAPs altered the staining of hair implants

Histological measurements of hair implant’s cross-sectional thicknesses revealed no significant differences (p = 0.478) among groups at 100 ± 6 microns on average. Although they have similar sizes and rounded morphologies, their MT staining appearances were generally different. REG and BLH appeared mostly yellow to yellowish, while M-KAP and L-KAP were more reddish with an occasional blue coloration (**Fig. 6E**), indicating that KAPs affected and/or interfered with aniline blue and Biebrich scarlet (red) dyes’ [29] protein interactions. Keratins are intermediate filament intracellular cytoskeletal proteins in the cytoplasm; thus, they expectedly stain red in MT as we observed in a previous publication [7] and are pronounced in M-KAP and L-KAP (**Fig. *5*B** and **Fig. 6E**). This evidence further supports an indirect correlation with decreased KAPs content.

#### Hairs with depleted KAPs seemingly had fewer immune cells on their surfaces

A high-magnification investigation of histological slices, with attention to the interface between implant and host tissue (**Fig. 7**) revealed two trends. First, REG and mostly BLH strands, which appeared yellowish (and contained high KAPs), also contained a higher number of immune cells (**Fig. 7**: REG and BLH), that are morphologically likely to be macrophages (Mφ) and mast cells, on their surfaces. These immune cells (red) were present individually or in multi-cellular clusters without any surrounding collagen ECM (blue in MT) on the biomaterial surface (hair cuticle), with more aggregation qualitatively seen in BLH. Despite the clustering of macrophages in some areas, no local chronic inflammatory macrophagic foreign body giant cells (FBGCs) forms were detected. Some of the immune cells close to the implant’s surface appeared to be connective tissue-resident granulocytes, the mast cells, as they contain distinct granules [30]. These granulocytes are not likely neutrophils, which are associated with acute inflammation, due to the length of time and absence of polymorphonuclear features [31]. Second, some BLH, M-KAP, and L-KAP fibers with reddish coloration (depleted KAPs) had deposited collagen ECM with fibroblast cells instead and had relatively fewer associated immune cells. Additionally, fibroblasts in M-KAP showed a more spread shape compared to spindle forms in L-KAP samples (**Fig. 7**: M-KAP and L-KAP), suggesting activated versus quiescent states, respectively [32].

**Fig. 7.**
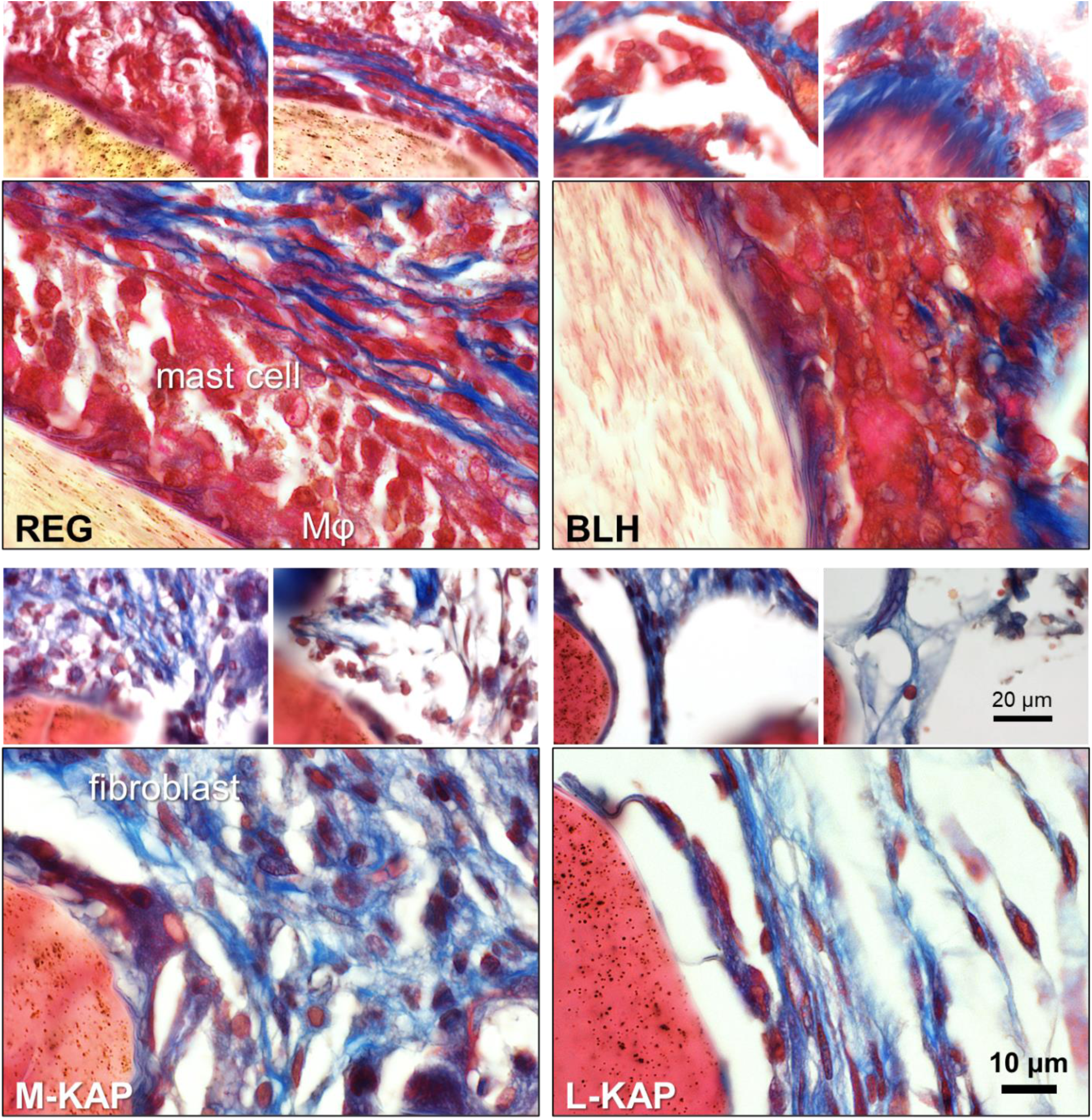
Hair biomaterial implants and their up-close interfacial interaction to the foreign body response cells and ECM components. Three representative images per group: 2 at lower magnification and 1 (boxed main) at higher showing clusters of immune cells (red) of mostly Mφ = macrophages and mast cells, and fibroblasts within the collagen ECM (blue).

Blood vessels associated with the FBR or the neoangiogenic vascular networks in the bulk of the fibrous capsule were found to be statistically the same (p = 0.526) across groups (**Fig. 8**), at 13 ± 5 vessels per mm^2^. Their small sizes and uni-layered shapes indicated them as endothelial tissues and capillaries (**Fig. 8**: REG) with an absence of smooth muscles in the FBR granulation tissue. Endothelial cells and their supporting pericytes and occasional macrophages were seen. Also, in the FBR bulk and outer surface, the collagen ECM appeared similar in morphologies across implant groups under high magnification (**Fig. 8**) with comparable densities (also confirmed in **Fig. 6D** image analysis). Occasional fibroblasts were detected in regions of dense bundles, with more spindle and stretched morphologies in the L-KAP samples.

**Fig. 8.**
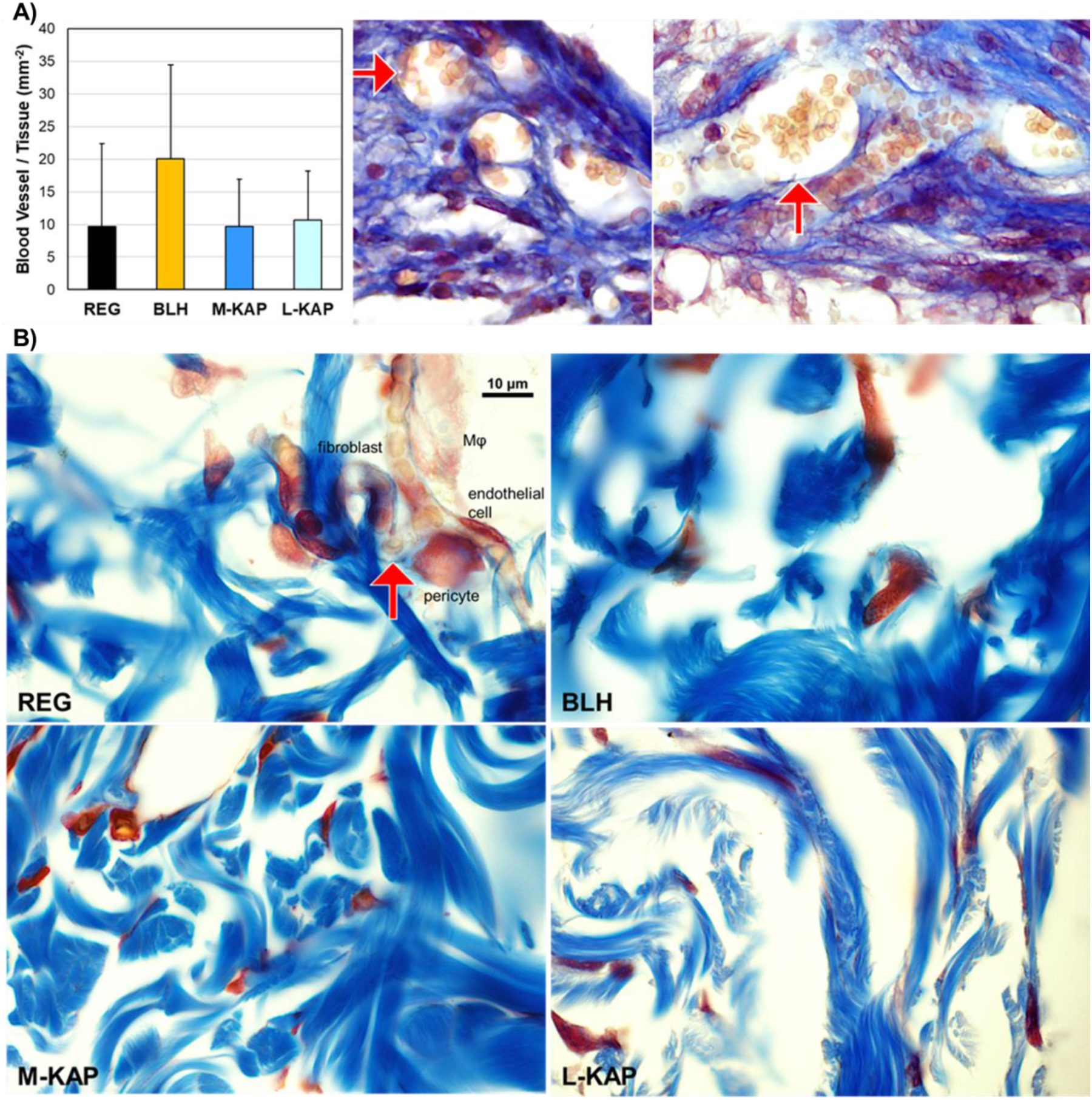
**A)** Blood vessels in the FBR fibrous tissue against hair implants (examples as arrows in the L-KAP group, with visible red blood cell contents). **B)** Collagen fiber bundles of the fibrous capsule against the hair implants. Branching capillaries (arrow) with endothelial cells, pericyte, and red blood cells delivered to the fibroblast cell of the fibrous capsule can be seen in REG. A Mφ = macrophage is also observed right next to the blood vessel.

The summary of local implantation effects is found in **Table 2**, where the H-KAPs (REG and BLH) scored to have a slight reaction, while M-KAP and L-KAP deemed to incite minimal or no reaction.

**Table 2.** Semiquantitative scoring of local effects after implantation.

| Hair groups | Score |  |  | Implantation effects |
| --- | --- | --- | --- | --- |
|  | AVE | STD | Reference = 2.9 subtracted from AVE |  |
| REG | 6.0 | 4.4 | 3.1 | slight |
| BLH | 7.8 | 3.6 | 4.9 | slight |
| M-KAP | 4.8 | 3.7 | 1.9 | no reaction or minimal |
| L-KAP | 3.5 | 1.0 | 0.6 | no reaction or minimal |

### 3.4 Summary of findings

Quantitative and semi-quantitative/qualitative analysis of human hair properties and their biocompatibility responses are summarized in the tables below. Although all hair biomaterials passed the safety *in vitro* cytotoxicity and incited only a thin fibrous capsule response, based on microscopic cell and ECM analysis, the concluding ranking of the most to “least” biocompatible is as follows: L-KAP, M-KAP, REG, and BLH.

## 4. Discussion

### Rationale behind the good biocompatibility of hair biomaterials

Residual hair biomaterials, including untreated hair, were all expectedly determined to be safe and biocompatible (**Table 3**). The subcutaneous fibrous capsule response (**Fig. 6**) was thin (with 28% mean) compared to a much thicker reaction in pacemaker implants in the mm range at ∼ 100% of the dermis thickness [24]. The paper’s objective outcome correlated with numerous previous studies on extracted and reassembled keratins and KAPs or the broad term “keratin biomaterials,” with or without other accepted polymeric biomaterials, showing good *in vitro* and *in vivo* biocompatibilities [10, 12, 33–35]. In the absence of the hair follicle which is the antigenic region [36] containing intact living cells, the terminal hair shaft or fiber is a non-immunogenic non-living biomaterial primarily due to its highly-conserved protein molecular structures across species [3, 37, 38]. Whenever immune cells from the same (allogeneic) or from different (xenogeneic: human hair into mouse sWAT) species get in contact with the hair implant, they are already “familiar” with keratins and KAPs, hence the foreign body response/reaction (FBR) is minimal or toned-down. The presence and intactness (for the most part) of the cuticle (**Fig. 2** and **Fig. 3**) prevented hair implant’s degradation and absorption, versus samples with cuticle intentionally removed and oxidatively-modified extracts that were almost completely resorbed at the 2-week time point [7]. This cuticular structure also largely influences the host immune response as its surface directly interacts (**Fig. 7**) with the host cells and extracellular matrix (ECM).

**Table 3.** Biocompatibility responses to hair samples.

| Hair groups | *In vitro*, fibroblast culture | *In vitro*, mouse subcutaneous fibrous capsule foreign body response ISO 10993-6 local effects after implantation |  |  |  |  |  | Overall relative biocompatibility |
| --- | --- | --- | --- | --- | --- | --- | --- | --- |
|  | ISO 10993-5 cell viability to extracts | Wall thickness vs. dermis | Tissue area per implant | Cells per tissue | Collagen density (mm<sup>-2</sup>) | Blood vessel density (mm<sup>-2</sup>) | Implant interface and local effects |  |
|  | Fig. 4B | Fig. 6A | Fig. 6B | Fig. 6C | Fig. 6D | Fig. 8A | Fig. 7 |  |
| REG | 98% ± 26%, pass | 32% ± 17%, thin | 10.1 ± 5.8 | 44% ± 4% | 1.8 ± 0.1 | 10 ± 13 | immune cells, slight | **good** |
| BLH | 84% ± 24%, pass | 43% ± 12%, thin | 15.8 ± 10.7 | 48% ± 3% | 1.9 ± 0.1 | 20 ± 14 | immune cells, slight | **decent** |
| M-KAP | 101% ± 7%, pass | 19% ± 8%, very thin | 1.9 ± 1.2 | 42% ± 4% | 1.7 ± 0.1 | 10 ± 7 | fibrous tissue, minimal | **better** |
| L-KAP | 106% ± 24%, pass | 19% ± 5%, very thin | 2.1 ± 0.9 | 38% ± 7% | 1.8 ± 0.1 | 11 ± 7 | fibrous tissue, minimal | **best** |

The cuticle’s outermost layer is the epicuticle or the fiber cuticle surface membrane containing the abundant 18-methyleicosanoic acid (18-MEA) lipid [39], which was removed or delipidized in this work for hydrophilicity and increased water penetration. Underneath, there are 6 to 10 flat and overlapping scaly layers or cuticular cells (we observed 5-7 in REG), each bound and bordered by a cell membrane complex (CMC) with an inside A-layer, exocuticle, and endocuticle. These are similar in their structure and proteomics to that of skin corneocytes or stratum corneum of the epidermis [3, 40–44]. Biochemically, the cuticle is also mostly composed of keratins and KAPs, with some differences in specific type and abundance compared to those in the cortex. The elevated keratins (Ks) in the cuticle region are known to be K1, K2, K10, K32, K40, K82; while those lower are K31, K33A, K33B, K38, K39, K83, K85, K86. The following groups of KAPs are mostly localized in the cuticle: KAP5, KAP10, KAP12, KAP17, and KAP19 [40, 41]. We infer that the identities, structural organization, and stability of these proteins are predominantly responsible for the relatively good FBR to hair biomaterials.

### Release mechanism of cortical KAPs

The decrease of KAPs in residual hairs (**Table 1**) from the cortex (**Fig. 3**) was correlated with the favorability of the biocompatibility trend (**Table 2** and **Table 3**), specifically the further decrease in size of the fibrous capsule and lower number of immune cells interacting with the implant’s surface. The implication is that cortical KAPs of REG and BLH hair leaches out, which causes an increase in cellular immune migration and activation.

Keratins and KAPs contain higher numbers of cysteine residues (with sulfur) compared to other proteins [45]; but among themselves, keratins are considered LS = low sulfur / low cysteine (Cys) at 3%-6.5% by moles [46]. In the hair cortex, keratins make up the intermediate filaments (IFs), but the globular and more polar KAPs of ∼ 2 nm in size [47] fill in the spaces around it and within the macrofibril bundles of the cortical cells. They are also extensively crosslinked via transglutamination and mostly disulfide bonding of cysteines [3, 44]. Out of the 89 different KAPs in human hair, those found in the cortex grouped into three families are: KAPs 1-3, 13, 15-16, and 23 as HS = high sulfur (11%-28% Cys), KAPs 4 and 9 as UHS = ultrahigh sulfur (29.5%-38% Cys), and KAPs 6-7 and 20-22 as HGT = high glycine and tyrosine (0-20% Cys), with MWs ranging from 8-31 kDa [19, 41, 48]. A possible explanation is that polar KAPs of the HGT family, particularly with low cysteines and low MW such as KAPs 6.1, 6.2, 7.1, 20.1, 20.2, 20.3, 22.1, and 22.2 (with 7%-15% Cys at 5-9 kDa) [49], detach more easily through absorbed water interaction, due to less crosslinking and smaller size (more mobile), respectively. They can diffuse out of the exposed edges of the cuticle since each scale is directly communicating or contacting the inner cortex, where water flowing in and out of the hair fiber is also controlled [3].

Aside from their main structural roles, intracellular KAPs, especially those in the epithelia, additionally assume non-mechanical and signaling functions alongside with IF keratins to facilitate normal skin maintenance [38, 50]. KAP5.5 was discovered to regulate the cytoskeletal function of cancer cells for motility and vascular invasion [51]. However, no information was publicly available on extracellular KAPs, such as those that are possibly released or secreted outside of cells, and their effects and actions on immune cells. Compared to keratins and other IF proteins, there is a larger variety of KAPs and IF-associated proteins. With higher variability comes less conservation among species [48], suggesting elevated immunogenicity of IF associated proteins (including KAPs) compared to IFs themselves [52].

Experimental groups with depleted KAPs imply that they contain purified keratins, in which the FBR including on their surfaces, are associated with less immune cells but instead with more deposition of collagen ECM (**Fig. 7**: M-KAP and L-KAP). Self-assembled extracted keratins (with higher purity, but still with melanin) are compatible with cultured primary fibroblasts and implicated with increase in collagen deposition *in vivo* [53, 54].

### Oxidation and associated cellular immune responses

Although satisfactory in biocompatibility, bleached hair was the least-favorable among the samples (**Table 3**). Hydrogen peroxide and persulfate (active ingredients of the lightener) permeate the hair and oxidatively decompose melanin granules under alkaline pH, rendering them water-soluble for removal during rinsing [55, 56]. Nevertheless, the bleaching process leads to the oxidation of amino acids of keratins and KAPs in the cuticle and cortex [1]. These modifications can recruit and activate local macrophages and mast cells via cell-surface pattern recognition receptors which detect non-self or damaged structures, then respond via phagocytosis and communication with fibroblasts for fibrous encapsulation [57–60]. Moreover, bleaching can also facilitate the removal of KAPs, as we observed in some BLH fibers appearing reddish in trichrome. In these instances, there were more fibrous tissues and less immune cells (**Fig. 7**: BLH). Perhaps, a more extensive rinsing is needed to remove excess of soluble oxidants, so the removal of KAPs outweigh the negative effects of native protein oxidation.

## 5. Conclusions

The insoluble residual hair strands that underwent treatment with the removal of melanin granules or KAPs in the cortex, as well as the untreated dark hair control, all displayed satisfactory *in vitro* and *in vivo* biocompatibilities according to ISO 10993 standards; therefore, passing the preliminary preclinical FDA requirements for safety of medical devices and/or biologics. Interestingly, human hair containing lower levels of KAPs and inferred to have higher purity levels of keratins with preserved intermediate filament, macrofibrillar, and cuticular structures, achieved the best result, with minimal immune cell presence on its surface, as well as thin fibrous encapsulation. Future steps towards clinical translation require testing on the effectiveness of the KAPs-depleted residual hair for a targeted tissue engineering and regenerative medicine application.

## CRediT authorship contribution statement

**Allison Meer**: Investigation, Formal analysis, Data curation, Writing – review & editing. **Aidan Mathews**: Investigation, Formal analysis, Data curation. **Mariana Cabral**: Investigation, Writing – review & editing. **Andrew Tarabokija**: Investigation. **Evan Carroll**: Investigation, Formal analysis. **Henna Chaudhry**: Investigation, Formal analysis, Writing – review & editing. **Michelle Paszek**: Investigation. **Nancy Radecker**: Investigation. **Thomas Palaia**: Investigation. **Roche C. de Guzman**: Conceptualization, Methodology, Resources, Investigation, Formal analysis, Visualization, Data curation, Writing – original draft, Writing – review & editing.

## Data availability

Data will be made available on request.

## Acknowledgements

We would like to thank Fusion Beauty Salon, Stylush Salon and Laser, Fresh Cuts, and Hair Designers for sources of human hair samples, Jason Williams for SEM assistance, and Atara Israel, Phoebe Christake, Hazel de Guzman, Jayda Lewis, and Alice Shvartsberg for help with samples processing, data analysis, and manuscript review.

